# V-TRACE: End-to-end temporal inference and annotation of animal behaviors from video

**DOI:** 10.64898/2026.04.14.718392

**Authors:** Kunming Shi, Guang-Wei Zhang, Can Tao, Zike Wang, Sonia K. Zhang, Huizhong W. Tao, Li I. Zhang

## Abstract

Quantitative analysis of animal behavior is fundamental to neuroscience and ethology but remains constrained by the scalability, subjectivity, and limited reproducibility of manual annotation. Many automated approaches infer behavior through intermediate representations such as pose trajectories, whereas direct video modeling retains motion, appearance, and context, but its accuracy and efficiency remain incompletely established. Here we introduce V-TRACE (<u>V</u>ideo-based <u>T</u>emporal <u>R</u>ecognition and <u>A</u>nnotation of <u>C</u>ontinuous <u>E</u>thograms of Animal Behavior), an end-to-end platform with a graphical user interface for behavioral detection and annotation. V-TRACE combines transformer-based video encoders with a multi-scale temporal module to model behavioral dynamics across a broad range of durations. Its modular design supports interchangeable encoder architectures within a common detection pipeline to produce frame-resolved behavioral identities and derive temporal boundaries from continuous recordings. Across datasets spanning species and experimental contexts, V-TRACE demonstrates high accuracy and high-throughput inference, enabling scalable, efficient, and context-aware animal behavior analysis directly from video.

## Introduction

Animal behavior unfolds as temporally structured actions embedded within continuous visual streams, where both movement dynamics and surrounding context contribute to behavioral identity^1–4^. Quantitative analysis of such behavior requires identifying and localizing discrete behavioral episodes over time, a process that remains constrained by the scalability, subjectivity, and limited reproducibility of manual annotation^1-6^. Despite the increasing availability of large-scale behavioral video recordings, scalable automated methods for behavioral annotation remain limited.

Pose-based methods have substantially advanced behavioral quantification, providing interpretable and precise descriptions of movement across diverse experimental contexts (DeepLabCut^7^, SLEAP^8^, LEAP^9^ and DANNCE^10^). The keypoint representation is itself an advantage for many questions because it supports biomechanical analysis, inter-individual distance, group heading and other measurements that cannot be obtained from behavior labels alone. Inferring behavioral states from pose trajectories involves an additional analytical stage applied to keypoint data (Keypoint-MoSeq^11^, MARS^12^ and SimBA^13^), such that behavior analysis involves first estimating pose throughout the recording and then inferring behavior from the resulting trajectories.

Recent advances in video representation learning offer the possibility of mapping video to temporally resolved behavioral labels within a single end-to-end pipeline, while retaining visual information beyond landmark coordinates, including appearance and environmental context that may contribute to behavioral identity^14-16^. These approaches are beginning to be explored in animal behavior analysis^17,18^. Transformer architectures and large-scale self-supervised learning have produced general visual representations^19-21^ and temporal video models^22-25^. In human action recognition, such models enable accurate action classification (VideoMAE^24,25^) and temporal detection^22,23^. Yet recognizing animal behavior presents distinct challenges relative to human action recognition: animal-behavior datasets, although increasing in availability and recording duration, remain substantially smaller and less diverse than the large human-video datasets used for general-purpose representation learning^26,27^; behavioral bouts can be sparse and highly imbalanced and can span widely varying durations; and recordings are typically continuous rather than pre-segmented into discrete clips^1,4^. Direct end-to-end temporal inference of animal behavior from continuous video therefore remains less established, particularly across diverse species and recording conditions.

Here we introduce V-TRACE (Video-based Temporal Recognition and Annotation of Continuous Ethograms of Animal Behavior), an end-to-end platform for temporal inference and annotation of animal behaviors from video. V-TRACE is implemented as a modular framework in which video encoder backbones, temporal modeling heads and adaptation strategies can be interchanged within a common video-to-behavior pipeline. Systematic comparison of these components identified a compact VideoMAEv2 ViT-B model distilled from a V-JEPA2 teacher^28^ as the default configuration, providing a favorable balance of behavioral accuracy, training and inference efficiency, and computational requirements. Across diverse laboratory and naturalistic datasets, V-TRACE produces temporally resolved behavioral labels across species and recording conditions within an integrated annotation, training, inference and correction workflow.

## Results

### V-TRACE performs end-to-end temporal inference of animal behavior from video

We developed V-TRACE as a modular platform for inferring temporally structured animal behaviors directly from continuous video recordings (**Fig. 1**). Human annotators use the graphical interface to mark behavioral identities and their start and stop times on the video timeline, generating temporally localized labels that are paired with the corresponding video frames for model training (**Fig. 1a**).

**Figure 1.**
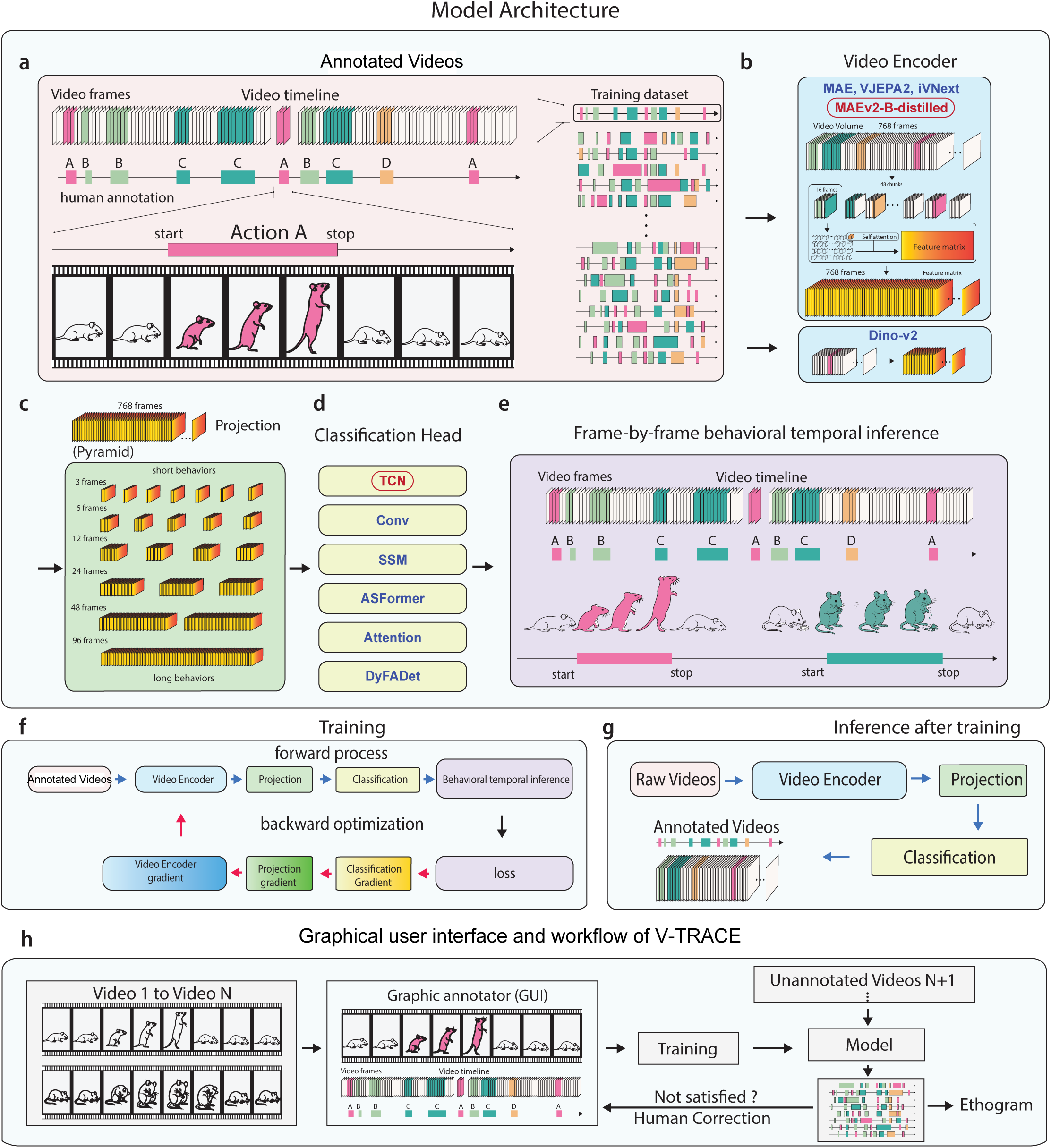
V-TRACE architecture and workflow for modular, frame-resolved behavioral inference. **a,** Annotated video training data. Raw videos are manually annotated with action onset and offset boundaries to generate temporal action sequences used as supervision during training; annotated clips are pooled into the training dataset (right). **b,** Interchangeable visual backbones. The 768-frame input is transformed into a temporally resolved feature matrix. For video encoders operating on 16-frame inputs, each video volume is divided into 48 non-overlapping temporal chunks, which are embedded and processed through self-attention layers. V-TRACE supports pretrained backbones from the MAE/VideoMAE, V-JEPA2, iVNext and DINOv2 families. The distilled MAEv2-B backbone, highlighted in red, is the default backbone used in this study. **c,** Projection into a multi-scale temporal pyramid. Encoder features are projected and represented at six temporal resolutions (3, 6, 12, 24, 48 and 96 frames), capturing behavioral events spanning short to long durations. **d,** Classification head. The temporal-modeling head is modular; six interchangeable classification-head implementations are shown: TCN, Conv, SSM, ASFormer, Attention and DyFADet. The TCN head, highlighted in red, is the default temporal head used in this study. **e,** Frame-by-frame behavioral temporal inference. V-TRACE assigns a behavioral class to every frame of the video timeline; contiguous runs of the same class define the start and stop boundaries of each behavioral bout. **f,** Training pipeline. Annotated videos pass through the video encoder, projection module and classification head in a forward pass; the loss is backpropagated through the trainable adapters, temporal projection and classification head to optimize model parameters. **g,** Inference after training. Raw videos are processed by the trained network to generate frame-wise behavioral annotations without additional manual labeling or pose estimation. **h,** Graphical user interface and end-to-end workflow of V-TRACE. Videos 1–N are labeled in the graphical annotator to produce frame-level annotations; the annotated set trains a model, which is applied to unannotated videos N+1 onward to generate ethograms. Predictions can be inspected and corrected by the user, and the corrected annotations can be fed back as additional training data for human-in-the-loop refinement.

V-TRACE encodes successive video chunks using an interchangeable video encoder, or backbone, and concatenates the resulting features along the temporal axis (**Fig. 1b**). The platform currently supports pretrained backbones from the MAE/VideoMAE^24,25^, V-JEPA2^28^, iVNext and DINOv2^21^ families within the same video-to-behavior pipeline. The encoded features are projected into a multi-scale temporal pyramid that represents behavioral information across progressively coarser temporal resolutions, enabling the model to accommodate behaviors ranging from brief movements to extended behavioral states (**Fig. 1c**). An interchangeable temporal classification head, such as TCN, Conv, SSM, ASFormer, Attention, DyFADet or TriDet (see **Methods**), then maps this multi-scale representation to frame-resolved predictions over all annotated behaviors and background (**Fig. 1d,e**). Unless otherwise specified, the experiments in this study used the default configuration highlighted in red in **Fig. 1b,d**: the distilled MAEv2-B encoder paired with the TCN classification head. The resulting output is an ethogram of behavioral identity over time; V-TRACE itself does not predict spatial masks, bounding boxes, keypoints or animal identity.

The same platform supports the complete annotation-to-inference workflow (**Fig. 1f-h**). V-TRACE is installed locally, and users launch its graphical interface on their computer. Users can load and annotate video, launch model training, apply the trained model to unannotated recordings, inspect and correct its predictions, and export the resulting ethogram. Corrected predictions can be retained as additional annotations, enabling an iterative human-in-the-loop workflow in which an initial model assists the annotation of subsequent recordings. V-TRACE produces one behavioral label stream for each video. V-TRACE was implemented so that video encoders, temporal heads and adaptation strategies can be evaluated within a common video-to-behavior pipeline. This modular design separates the integrated user workflow from the choice of model configuration and enables systematic comparison of accuracy, training cost, inference throughput and computational requirements, as examined below.

### Accurate and robust temporal detection of spontaneous mouse behaviors

We first evaluated the default V-TRACE configuration, the distilled MAEv2-B encoder paired with a TCN head, on recordings of single freely behaving mice annotated for self-grooming, rearing, drinking and eating (**Fig. 2a**). The annotated dataset comprised 505 clips from 13 mice across 34 recordings. Across these clips, bouts of each behavior occurred at different times and for different durations, interspersed with extensive unlabeled intervals (**Fig. 2b**). At the frame level, 70.7% of the annotated data carried no behavioral label. Among labeled behavioral frames, rearing was most frequent (49.9%), followed by eating (23.0%), self-grooming (16.5%) and drinking (10.6%) (**Fig. 2c**).

**Figure 2.**
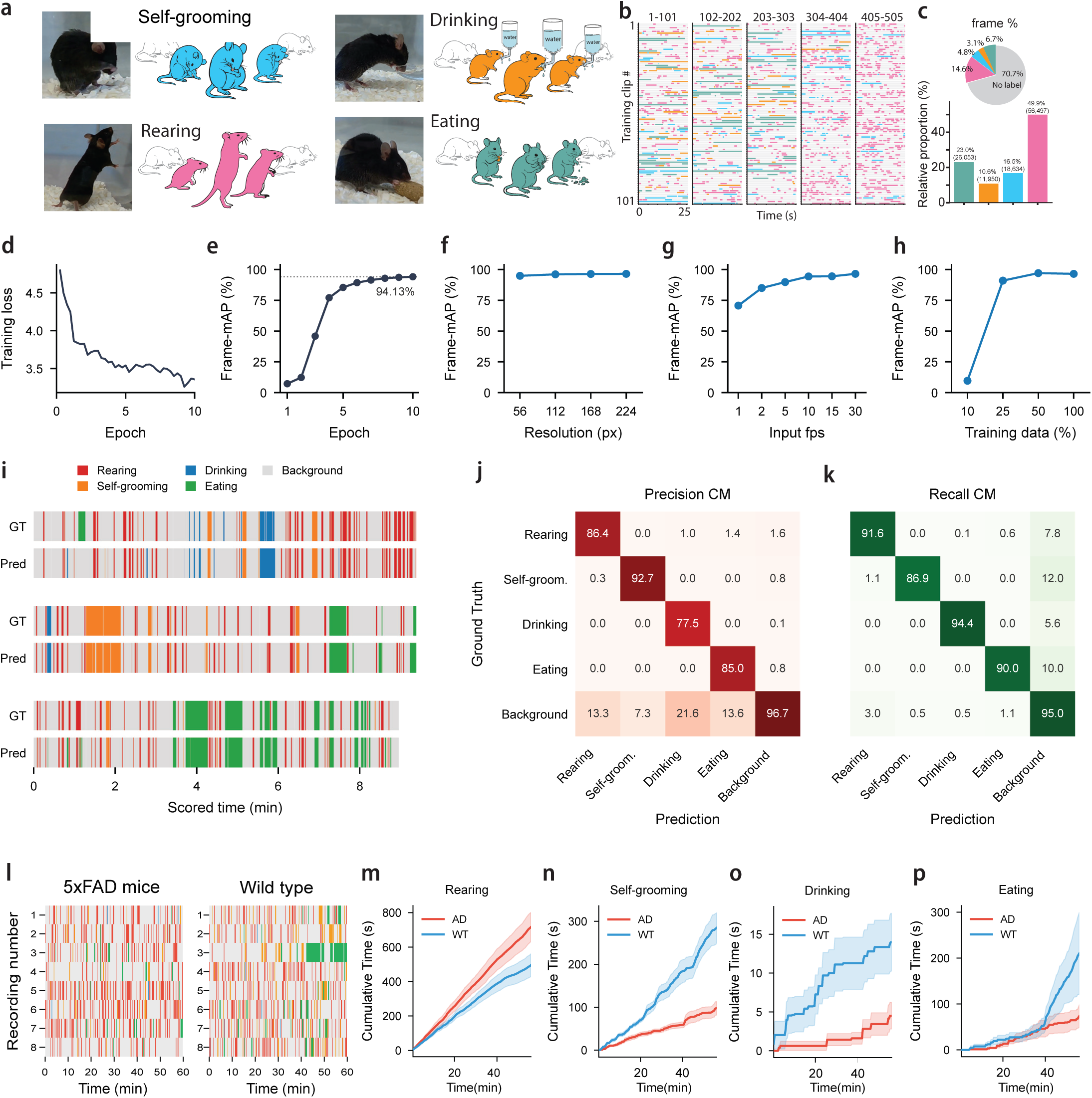
V-TRACE achieves accurate frame-wise detection of mouse home-cage behaviors. **a,** Representative video frames and schematics of the four annotated home-cage behaviors: self-grooming (blue), drinking (orange), rearing (pink) and eating (teal). Schematics show the behavioral bout (colored) flanked by non-behavior frames (grey outlines). **b,** Visualization of the 505-clip annotated dataset displayed in five blocks. Each row is one 25-s annotated clip; colored segments denote annotated behavioral bouts. **c,** Behavioral composition of the annotated dataset. Pie chart, per-frame proportions: rearing 14.6%, eating 6.7%, self-grooming 4.8%, drinking 3.1% and unlabeled intervals (“no label”) 70.7%. Bar plot, relative proportions among labeled frames with absolute frame counts: rearing 49.9% (56,497), eating 23.0% (26,053), self-grooming 16.5% (18,634) and drinking 10.6% (11,950). **d,** Training loss across 10 epochs. **e,** Held-out test performance across epochs. Frame-level mean average precision (frame-mAP) reached 94.13% at the final epoch. **f–h,** Robustness analyses showing frame-mAP as a function of input resolution (f; 56, 112, 168 and 224 px), input frame rate (g; 1, 2, 5, 10, 15 and 30 fps) and the fraction of the training portion used (h; 10, 25, 50 and 100%). **i,** Comparison of ground-truth (GT) and V-TRACE-predicted (Pred) behavioral timelines for three representative held-out test clips, showing temporal localization of behavioral bouts (per-clip frame accuracy 99.02%, 95.77% and 92.75%). **j, k,** Precision (j) and recall (k) confusion matrices for rearing, self-grooming, drinking, eating and background classes; values are percentages. **l,** Behavioral timelines across 60-min recordings from 5×FAD and wild-type (WT) mice (n = 3 mice per group; eight recording sessions per group), with each row representing one recording session. **m–p,** Cumulative time spent in rearing (m), self-grooming (n), drinking (o) and eating (p) in 5×FAD (AD) and WT mice, plotted against time (min). Shaded regions indicate mean ± s.e.m. *p* values were calculated using two-sided Mann–Whitney U tests: rearing *p* = 0.038, self-grooming *p* = 3.1 × 10⁻^4^, drinking *p* = 0.153 and eating *p* = 0.130.

Training loss decreased over the fixed optimization schedule, while frame-mAP on the held-out test portion rose rapidly and reached 94.13% at the last epoch (**Fig. 2d,e**; see **Methods**).

V-TRACE was robust to substantial changes in spatial resolution, with little variation in frame-mAP across the tested range of 56-224 pixels (**Fig. 2f**). Performance was also largely maintained at frame rates of 5-30 frames per second (fps) but declined as temporal sampling was reduced to 2 fps and deteriorated markedly at 1 fps (**Fig. 2g**). Reducing the training portion to 25% retained high performance, whereas using only 10% produced a pronounced loss of accuracy (**Fig. 2h**; see **Methods**). On representative held-out test clips, predicted and manually annotated behavioral sequences were closely aligned, with the three examples reaching frame accuracies of 99.02%, 95.77% and 92.75% (**Fig. 2i**). The precision confusion matrix (CM) showed relatively high class-specific precision (**Fig. 2j**). The recall matrix yielded recalls of 91.5% for rearing, 86.9% for self-grooming, 94.4% for drinking and 90.0% for eating, with most missed behavioral frames assigned to background rather than to another behavior (**Fig. 2k**).

We next used V-TRACE-generated ethograms to compare 5×FAD Alzheimer’s disease model mice^29^ with wild-type controls; the session-level timelines, arranged by group, revealed visibly different allocations of rearing, self-grooming, drinking and eating across the recording period (**Fig. 2l**). Cumulative behavioral time showed increased rearing in 5×FAD mice (*p* = 0.038; **Fig. 2m**) and reduced self-grooming (*p* = 3.1 × 10⁻^4^; **Fig. 2n**), whereas drinking (*p* = 0.153; **Fig. 2o**) and eating (*p* = 0.130; **Fig. 2p**) did not differ significantly between groups. Rearing and self-grooming are species-typical behaviors used as indices of exploration and compulsivity, respectively^30^, making their altered expression relevant to the behavioral phenotype of 5×FAD mice^31^. These results show that frame-resolved V-TRACE predictions can be aggregated to quantify group-associated differences across the recorded behavioral sessions.

To assess robustness to a separate validation split, we also divided the 505 annotated clips into training, validation and test subsets. The validation-selected model achieved 96.3% frame-mAP on held-out test clips (**Extended Data Fig. 1a,b**). Because this analysis partitioned data at the clip level, it measures performance on held-out clips but does not establish generalization to unseen recordings or animals.

### V-TRACE achieves high accuracy on the CalMS21 benchmark

We next evaluated V-TRACE on the Caltech Mouse Social Interactions (CalMS21) benchmark^32^, which contains resident-intruder recordings annotated for attack, investigation and mount, with unannotated frames treated as background (**Fig. 3a**). Clip-level timelines showed that the three social behaviors were unevenly distributed across the training set and interspersed with extensive unlabeled periods (**Fig. 3b**). Background accounted for 62.8% of all frames (**Fig. 3c**); among labeled behavioral frames, investigation predominated (78.8%), followed by mount (14.3%) and attack (6.9%) (**Fig. 3d**).

**Figure 3.**
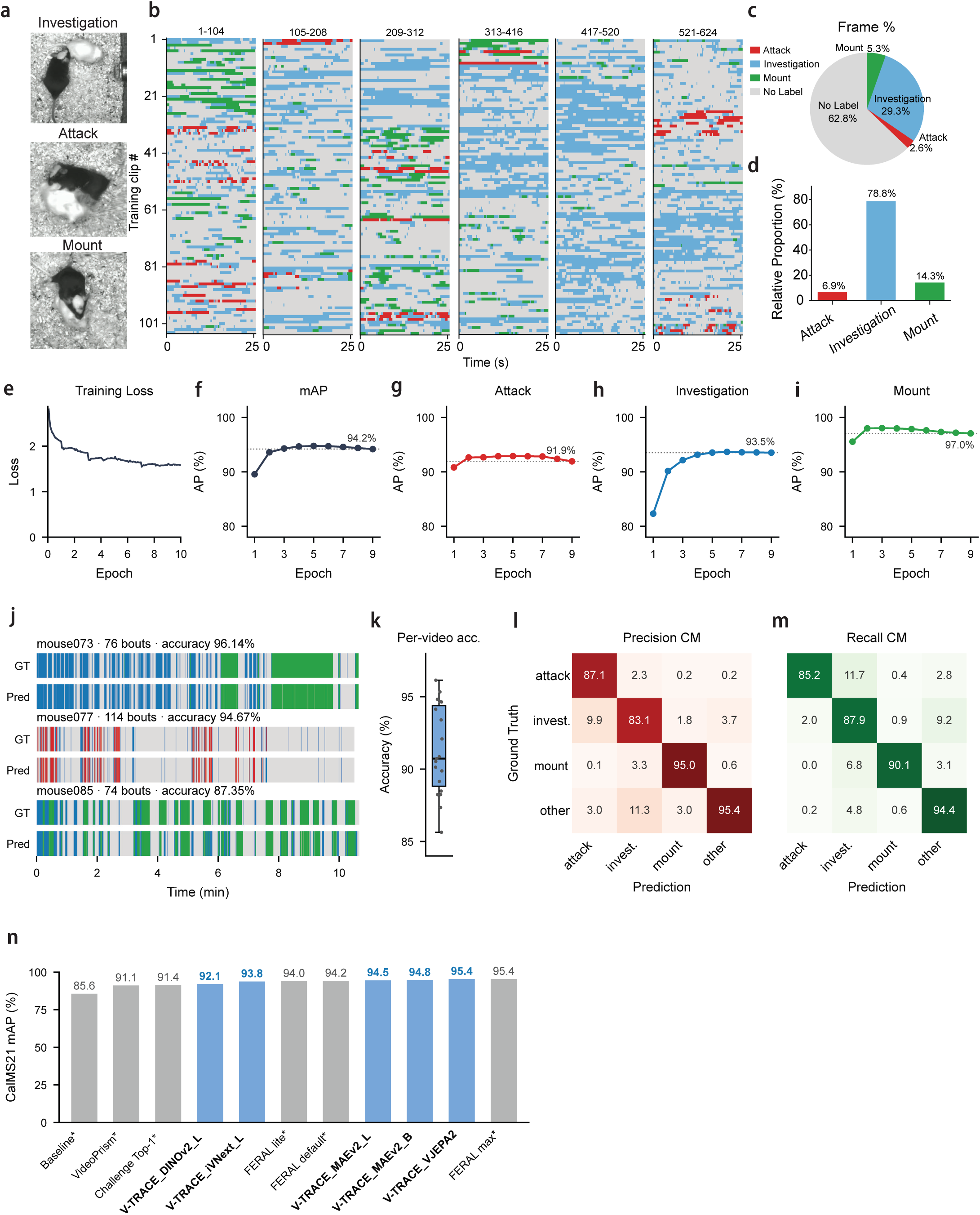
V-TRACE accurately localizes social behaviors in the CalMS21 mouse social-interaction benchmark. **a,** Representative frames illustrating investigation, attack and mount behaviors. **b,** Visualization of the training dataset displayed in six blocks (clips 1–104, 105–208, 209–312, 313–416, 417–520 and 521–624). Each row is one 25-s training clip; colored segments indicate annotated behavioral bouts and unlabeled intervals. **c,** Pie chart showing per-frame behavioral composition of the dataset: investigation 29.3%, mount 5.3%, attack 2.6% and no label 62.8%. **d,** Relative proportions among labeled frames: investigation 78.8%, mount 14.3% and attack 6.9%. **e,** Training loss across 10 epochs. **f,** Mean average precision (frame-mAP) during training, reaching 94.2%. **g–i,** Per-class AP curves for attack (g; 91.9%), investigation (h; 93.5%) and mount (i; 97.0%). **j,** Comparison of ground-truth (GT) and V-TRACE-predicted (Pred) behavioral timelines for three held-out test videos (per-video frame accuracy 96.14%, 94.67% and 87.35%). **k,** Distribution of per-video classification accuracy across test videos; box shows median and interquartile range, whiskers the full range and points individual videos. **l,m,** Precision (l) and recall (m) confusion matrices for attack, investigation, mount and other classes; values are percentages. **n,** CalMS21 frame-mAP comparison. Grey bars show values reported for published comparator methods: the pose-based challenge baseline (85.6%), VideoPrism (91.1%), the challenge Top-1 entry (91.4%), and FERAL lite (94.0%), default (94.2%) and max (95.4%). Blue bars show encoders evaluated within the V-TRACE framework: DINOv2-L (92.1%), iVNext-L (93.8%), MAEv2-L (94.5%), MAEv2-B (94.8%) and V-JEPA2 (95.4%). Asterisks denote values reported by the original studies rather than results from matched reruns.

Training loss decreased and stabilized over the fixed optimization schedule (**Fig. 3e**), while epochwise frame-mAP reached 94.2% (**Fig. 3f**). Class-specific AP remained high but differed across behaviors, reaching 91.9% for attack (**Fig. 3g**), 93.5% for investigation (**Fig. 3h**) and 97.0% for mount (**Fig. 3i**). Representative test-video ethograms showed close temporal agreement between predictions and annotations, with example frame accuracies of 96.14%, 94.67% and 87.35% (**Fig. 3j**). Per-video accuracy was generally high but varied across the official test recordings (**Fig. 3k**). Precision and recall confusion matrices showed that residual errors occurred chiefly between attack and investigation or between investigation and background, while mount remained comparatively well separated (**Fig. 3l,m**).

Using the official 70-video training and 19-video test partition, the default distilled MAEv2-B V-TRACE configuration achieved 94.8% frame-mAP, while V-TRACE with the larger V-JEPA2 encoder achieved 95.4% (**Fig. 3n**). Additional V-TRACE configurations based on DINOv2-L, iVNext-L and VideoMAEv2-L achieved 92.1%, 93.8% and 94.5%, respectively. Published results obtained under the same official benchmark include 85.6% for the pose-based challenge baseline^32^, 91.1% for VideoPrism^33^ and 91.4% for the challenge Top-1 entry^32^. FERAL^16^ uses V-JEPA2-family backbones across its lite, default and max configurations, which reported 94.0%, 94.2% and 95.4% frame-mAP, respectively (**Fig. 3n**). Because these systems were trained and selected by their original authors, the literature values should not be interpreted as matched reruns. The comparable performance of V-JEPA2-based models in the independently developed V-TRACE and FERAL pipelines is consistent with the usefulness of this encoder family for direct video-to-behavior inference.

Sensitivity to validation-set composition was evaluated separately using three mutually disjoint validation subsets drawn from the 70-video training partition. Validation-selected checkpoints achieved 95.23%, 94.82% and 94.75% frame-mAP on the same official test set (**Extended Data Fig. 1c,d**; Supplementary Table 1), indicating limited sensitivity to the selected validation partition. Because these runs used fewer training videos and a different checkpoint-selection rule than the main fixed-epoch analysis, the difference between the protocols cannot be attributed to either factor alone.

V-TRACE was additionally evaluated on the Mouse-Homecage and Mouse-Social benchmarks released with DeepEthogram^14^, using the corresponding partitioning protocol and F1 metrics (**Extended Data Fig. 2**; Supplementary Table 2). Across five video-level splits, V-TRACE achieved micro-F1 scores of 0.861 on Mouse-Homecage and 0.925 on Mouse-Social, compared with published DeepEthogram values of 0.760 and 0.920, respectively. On CalMS21, V-TRACE achieved a micro-F1 of 0.922, compared with 0.833 for the DeepEthogram pipeline evaluated under the corresponding protocol (**Extended Data Fig. 2c**).

### Cross-species behavioral detection in Drosophila and chimpanzee datasets

To evaluate V-TRACE across species and recording contexts, we first applied the method to Drosophila courtship videos^34^ annotated for circle, copulation and wing extension (**Fig. 4a**). Clip-level timelines revealed pronounced imbalance and substantial variation in behavioral composition across the training set (**Fig. 4b**). Copulation occupied 76.3% of all frames, compared with 1.9% for wing extension and 0.5% for circle, while 21.4% of frames were unlabeled (**Fig. 4c**). Among labeled behavioral frames, copulation therefore represented 97.0%, wing extension 2.4% and circle 0.6% (**Fig. 4d**).

**Figure 4.**
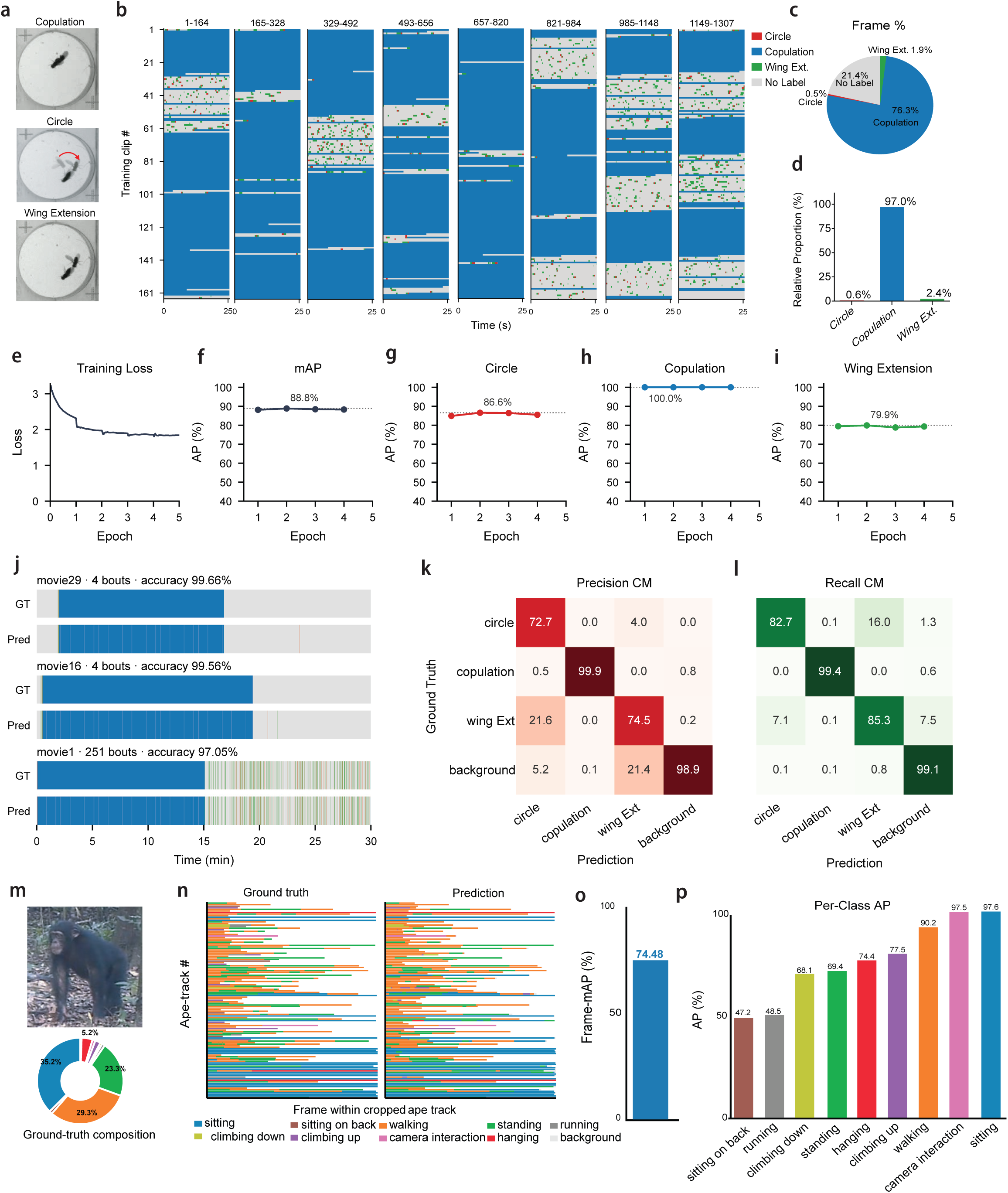
V-TRACE detects temporally resolved behaviors across Drosophila courtship and chimpanzee camera-trap datasets. **a,** Representative frames of Drosophila courtship behaviors: copulation, circle and wing extension. **b,** Visualization of the Drosophila training dataset displayed in eight blocks (clips 1–164, 165–328, 329–492, 493–656, 657–820, 821–984, 985–1148 and 1149–1307). Each row is one 25-s training clip; colored segments indicate circle, copulation, wing extension and unlabeled intervals. **c,** Pie chart of per-frame behavioral composition in the Drosophila dataset: copulation 76.3%, wing extension 1.9%, circle 0.5% and no label 21.4%. **d,** Relative proportions among labeled Drosophila behaviors: copulation 97.0%, wing extension 2.4% and circle 0.6%. **e,** Training loss across 5 epochs. **f,** Mean average precision (mAP) during training, reaching 88.8%. **g–i,** Per-class AP curves for circle (g; 86.6%), copulation (h; 100.0%) and wing extension (i; 79.9%). **j,** Comparison of ground-truth (GT) and V-TRACE-predicted (Pred) Drosophila behavioral timelines for three held-out movies (per-video frame accuracy 99.66%, 99.56% and 97.05%). **k, l,** Precision (k) and recall (l) confusion matrices for the Drosophila dataset; values are percentages. **m,** Representative frame from the chimpanzee PanAf500 dataset and ground-truth composition of the annotated behaviors: sitting 35.2%, walking 29.3%, standing 23.3%, hanging 5.2% and remaining classes (running, climbing up, climbing down, sitting on back and camera interaction). **n,** Ground-truth (left) and V-TRACE-predicted (right) behavioral timelines across cropped chimpanzee tracks; each row is one chimpanzee track. **o,** Overall frame-level mAP on the held-out PanAf500 test partition (74.48%). **p,** Per-class average precision for chimpanzee behaviors: sitting 97.6%, camera interaction 97.5%, walking 90.2%, climbing up 77.5%, hanging 74.4%, standing 69.4%, climbing down 68.1%, running 48.5% and sitting on back 47.2%.

Training loss decreased and stabilized within five epochs (**Fig. 4e**). In the best-performing fold shown in **Fig. 4**, V-TRACE achieved 88.8% frame-mAP (**Fig. 4f**), comprising AP values of 86.6% for circle (**Fig. 4g**), 100.0% for copulation (**Fig. 4h**) and 79.9% for wing extension (**Fig. 4i**). Representative timelines closely matched the annotated sequences, with example frame accuracies of 99.66%, 99.56% and 97.05% (**Fig. 4j**). The precision matrix indicated that the rare circle and wing-extension classes were more susceptible to contamination from other labels or background (**Fig. 4k**), whereas the recall matrix showed recalls of 82.7% for circle, 99.4% for copulation and 85.3% for wing extension (**Fig. 4l**).

We used video-level five-fold analysis to quantify sensitivity to the movies assigned to training, validation and test subsets. Held-out frame-mAP across the five folds was 85.89%, 88.18%, 83.59%, 88.77% and 88.83%, giving a mean of 87.05 ± 2.27% (s.d.; n = 5 folds; **Extended Data Fig. 1e-g**). The observed variation across folds provides a distribution of performance across alternative movie-level partitions and a more representative estimate than any single fold.

We next evaluated V-TRACE on PanAf500 camera-trap recordings^35^ of wild chimpanzees. The test data contained a heterogeneous behavior distribution dominated by sitting, walking and standing, together with several less frequent behaviors (**Fig. 4m**). Because several chimpanzees can appear in the same frame while performing different behaviors, a single whole-frame behavioral label is often not well defined. Following the PanAf workflow, chimpanzees were detected and tracked externally, and V-TRACE classified the subject-centered crop associated with each track; detection and tracking were preprocessing steps external to V-TRACE, and no pose or keypoint representation was used. Across ape tracks, predicted behavioral timelines reproduced much of the temporal structure of the ground-truth annotations while retaining visible class-specific errors (**Fig. 4n**). On the held-out PanAf500 test partition, this correspondence produced an overall frame-mAP of 74.48% (**Fig. 4o**). Per-class AP varied substantially, ranging from 47.2% for sitting on back and 48.5% for running to 97.5% for camera interaction and 97.6% for sitting (**Fig. 4p**).

Compared with whole-frame inputs, subject-centered cropping increased AP for six of nine behaviors, including gains of 56.8 points for camera interaction and 47.9 points for hanging, but reduced AP for sitting on back, running and standing (**Extended Data Fig. 3**). Thus, subject-centered cropping improved recognition for several behaviors, whereas preserving the surrounding scene remained beneficial for others. Together, these analyses show that V-TRACE can produce temporally resolved behavioral labels across mouse, fly and chimpanzee recordings. They also define an important boundary of the approach: when different animals perform different behaviors simultaneously, subject identity must be resolved before individual behavioral streams can be assigned. Subject-centered crops provide one such input, but the balance between subject localization and surrounding context remains behavior dependent. Performance also remains variable for sparsely represented behaviors in unconstrained footage.

### Modular model design balances accuracy and computational efficiency

The modular structure of V-TRACE enabled us to compare video encoders, temporal heads and adaptation strategies within a common behavioral-detection pipeline (**Fig. 5**). In a controlled comparison of seven fully fine-tuned encoders on CalMS21, frame-mAP ranged from 89.16% for VideoMAEv2 ViT-S to 93.76% for V-JEPA2 (**Extended Data Fig. 4**). Performance generally increased with model scale, and V-JEPA2 achieved the highest accuracy under this comparison, motivating its use as the teacher for cross-architecture distillation.

**Figure 5.**
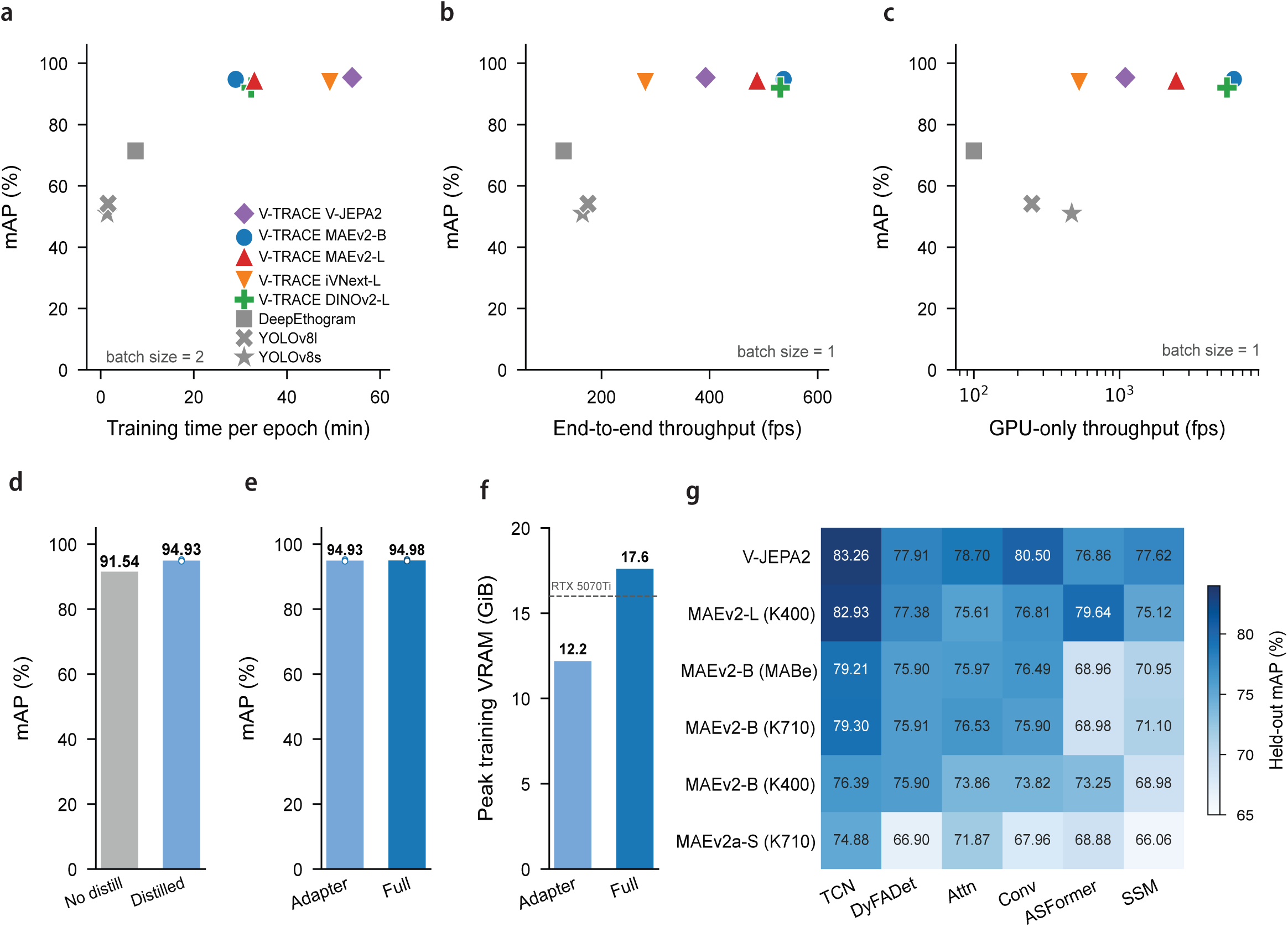
V-TRACE characterizes accuracy–efficiency trade-offs across video encoders, fine-tuning strategies and temporal heads. All comparisons were performed on CalMS21. **a–c,** Accuracy–cost trade-offs for five V-TRACE encoder configurations—V-JEPA2, MAEv2-B, MAEv2-L, iVNext-L and DINOv2-L—and three comparator methods—DeepEthogram, YOLOv8l and YOLOv8s. Colors and symbols are defined in a and used consistently across a–c. Frame-mAP is plotted against training time per epoch at batch size 2 (a; minutes), end-to-end inference throughput at batch size 1 (b; frames per second) and GPU-only inference throughput at batch size 1 (c; frames per second, logarithmic scale). **d,** Effect of encoder distillation on mAP (no distillation 91.54% vs distilled 94.93%). **e,** Effect of the fine-tuning strategy on mAP (adapter tuning 94.93% vs full fine-tuning 94.98%). **f,** Peak training VRAM for the two fine-tuning strategies (adapter 12.2 GiB vs full 17.6 GiB); dashed line marks the 16 GiB memory of an RTX 5070 Ti. **g,** mAP for all combinations of pretrained video encoder (rows: V-JEPA2, MAEv2-L (K400), MAEv2-B (MABe), MAEv2-B (K710), MAEv2-B (K400), MAEv2a-S (K710)) and temporal classification head (columns: TCN, DyFADet, Attn, Conv, ASFormer, SSM).

We next compared held-out accuracy with training time across V-TRACE configurations and matched DeepEthogram and YOLOv8 baselines^36^. All tested V-TRACE encoders achieved substantially higher mAP than these baselines, while the compact MAEv2-B configuration required the shortest training time among the V-TRACE models and retained near-top accuracy (**Fig. 5a**). End-to-end throughput, which includes video decoding and pipeline overhead, varied substantially among V-TRACE encoders; MAEv2-B provided the highest end-to-end throughput among the tested V-TRACE configurations while maintaining approximately 95% mAP (**Fig. 5b**). GPU-only throughput, which isolates model execution, was higher than end-to-end throughput for every configuration, confirming that decoding and other pipeline operations contribute appreciably to deployed runtime; MAEv2-B again combined the highest throughput with near-top accuracy (**Fig. 5c**). Together, these comparisons characterize practical trade-offs rather than establishing that one configuration is optimal for every application.

Cross-architecture distillation increased the held-out frame-mAP of the VideoMAEv2 ViT-B student from 91.54% to 94.93% without changing the student architecture or adding inference-time model components (**Fig. 5d**). Adapter-based fine-tuning achieved 94.93% frame-mAP, nearly matching the 94.98% obtained by full fine-tuning (**Fig. 5e**). It simultaneously reduced peak training GPU memory from 17.6 to 12.2 GiB, bringing the requirement below the capacity of a 16-GB consumer GPU (**Fig. 5f**). Thus, distillation transferred much of the accuracy of the larger teacher into the compact student, while adapter tuning reduced the memory required to train it. Six temporal-head families were evaluated across six encoder configurations (**Fig. 5g**). TCN provided consistently high mAP across the tested encoder configurations and was therefore selected as the default, although the relative differences among heads varied with the backbone. These results also show that temporal-head performance remains encoder dependent.

We further evaluated the complete default configuration on a 16-GB GeForce RTX 5070 Ti (**Extended Data Fig. 5**). On CalMS21, training performance remained robust across alternative validation partitions (**Extended Data Fig. 5a**). Across the four datasets, the deployed runs remained within the available GPU memory (**Extended Data Fig. 5b**) and achieved approximately 900-1,400 frames per second of end-to-end inference throughput (**Extended Data Fig. 5c**). These measurements show that the default model can be trained and applied without datacenter-class GPU hardware, although CPU-only inference and host-memory requirements were not evaluated.

Taken together, the systematic comparisons support the distilled, adapter-tuned MAEv2-B configuration as the default practical configuration because it provides a favorable balance of accuracy, training cost, inference throughput and GPU-memory requirements. Larger encoders remain viable when maximizing accuracy is the overriding priority, while alternative heads and backbones allow the platform to be adapted to different computational constraints.

## Discussion

In this study, we developed V-TRACE, an end-to-end and modular platform for frame-resolved temporal inference and annotation of animal behaviors directly from video. Across datasets spanning controlled laboratory assays and naturalistic camera-trap recordings, V-TRACE detected temporally structured behaviors in mice, Drosophila and wild chimpanzees, and its predicted ethograms supported the identification of behavioral differences in 5×FAD mice. On the CalMS21 benchmark, V-TRACE achieved high accuracy across multiple model configurations and remained robust to alternative validation partitions. Its modular design further enabled systematic comparison of interchangeable video encoders and temporal heads, while cross-architecture distillation and adapter-based fine-tuning transferred high accuracy to a compact model with reduced training-memory requirements. Together, these results support the use of V-TRACE as a practical framework for learning from expert-annotated video and generating temporally resolved behavioral measurements across diverse species, recording environments and experimental contexts.

Pose-estimation methods have substantially advanced automated behavioral analysis by providing interpretable keypoint trajectories for quantifying body configuration, joint motion, inter-individual distance and heading^7-9,37^. Inferring behavioral categories from these trajectories generally requires a subsequent analytical stage, and keypoint representations may not contain complete visual information, such as appearance, objects and environmental context, that may help to identify behaviors. Direct pixel-based approaches^14-16^ offer a complementary route by mapping visual and temporal information to behavioral labels without requiring pose as an intermediate representation. V-TRACE contributes to this emerging class by learning spatiotemporal representations that integrate motion, appearance and context across multiple temporal scales. The choice between direct-video and pose-based analysis therefore depends on the biological question rather than on a universal ordering of methods. Pose-based analysis is particularly appropriate when explicit kinematic variables are the primary readout, whereas direct-video inference is well suited to behaviors defined by temporally evolving visual patterns or contextual cues. Because both approaches produce frame-indexed outputs, V-TRACE-defined behavioral bouts can be aligned with pose-derived measurements when categorical behavior and kinematics are both required, although the current release does not automate this integration.

At a technical level, V-TRACE combines established advances in pretrained video representation learning and temporal action modeling within a modular, user-facing platform. Rather than tying behavioral inference to a single architecture, the platform supports interchangeable video encoders and temporal heads within a common annotation-to-inference workflow. The comparative analyses showed that high accuracy could be achieved across several encoder configurations and supported TCN as a practical default, while the relative performance of the alternative heads varied with the backbone. Cross-architecture distillation transferred much of the accuracy of the larger V-JEPA2 teacher to a compact MAEv2 ViT-B student without adding inference-time components, while adapter-based fine-tuning preserved this accuracy with lower training-memory requirements. Under the evaluated conditions, faster-than-real-time inference and deployment on a 16-GB consumer GPU support the feasibility of large-scale video inference using laboratory computing hardware. The graphical interface further connects annotation, training, inference and manual correction, allowing model predictions to assist the annotation of subsequent recordings.

Several limitations warrant consideration. V-TRACE relies on supervised annotations, and its performance depends on how well the training data represent the behavioral and visual variability encountered during inference. The cross-species analyses demonstrate that the same framework can be adapted across diverse experimental settings, but each dataset was trained separately using task-specific behavioral labels. In PanAf500, performance on less frequent behaviors reflected both limited training examples and the visual difficulty of distinguishing the behavior-relevant subject from the surrounding scene, indicating that class frequency alone does not determine accuracy. More broadly, applying V-TRACE to new experimental contexts will depend on both representative annotations and input preparation that preserves behavior-relevant visual information.

Taken together, V-TRACE illustrates how end-to-end spatiotemporal modeling can support structured behavioral quantification directly from video. Future extensions may incorporate improved video encoders and streamline adaptation to new datasets and experimental settings. By integrating contextual visual information with temporally resolved inference, V-TRACE provides a flexible approach for generating frame-resolved ethograms across diverse species and recording contexts.

## Acknowledgments

This work was supported by grants from the U.S. National Institutes of Health (DC008983, DC020887, NS132912 to L.I.Z. and EY019049, AG089756 to H.W.T.), Cure Alzheimer’s Fund to H.W.T.; Alzheimer Foundation Fellowship (AARF-23-1148428), Whitehall Foundation Research Grant (2026-08-034), VCU Convergence Award and 2026 Pilot Projects Award from Animal Models for the Social Dimensions of Health and Aging Research Network (NIH R24 AG065172) to G.W.Z.; and Brain & Behavior Research Foundation support to C.T. High-performance computing resources provided by the High Performance Research Computing (HPRC) core facility at Virginia Commonwealth University (https://hprc.vcu.edu) were used for conducting the research reported in this work. The Center for Advanced Research Computing (CARC) at the University of Southern California (https://carc.usc.edu) also provided computing resources for this study.

## Author Contributions

L.I.Z. and G.W.Z. conceived the study and supervised the project. K.S. and G.W.Z. adapted, implemented, and tested the V-TRACE model, designed and performed behavioral experiments, and conducted data analysis. S.K.Z., Z.W. and C.T. contributed to data analysis. G.W.Z., K.S., H.T. and L.I.Z. wrote the manuscript.

## Methods

### Animals

All experimental procedures in this study have been approved by the Institutional Animal Care and Use Committee (IACUC) of the University of Southern California. Male and female adult mice, C57BL/6J and 5×FAD, were used in this study. Mice were housed at 18–23 °C with 40–60% humidity in a 12-h light–dark cycle (6AM–6PM) with ad libitum access to food and water.

### Datasets

We evaluated V-TRACE across datasets spanning diverse species, recording conditions, and levels of behavioral organization. All datasets provide raw video paired with temporally annotated behavioral episodes. Training and evaluation details for each dataset are described below. Across all datasets, behavioral events were treated as non-overlapping and each frame was assigned at most one behavior label during annotation. For the single-mouse, CalMS21 and fly datasets, videos were supplied after resizing to the selected backbone’s required input resolution, without spatial cropping, background subtraction, tracking-based centering, detection or keypoint estimation; the subject-centered PanAf500 preprocessing is described below.

### Evaluation protocols

Datasets were evaluated under two partition schemes, a two-way train/test split and a three-way train/validation/test split, because the published baselines we compare against differ in whether they use a validation subset. In the two-way protocol, no validation subset was used and therefore no validation-based checkpoint selection was performed. Models were trained for a fixed dataset-specific schedule, and the last-epoch checkpoint was reported without selecting it according to test performance. The last epoch was not necessarily the highest-scoring epoch; in these runs, frame-mAP could peak a few epochs earlier and the last-epoch score was a few tenths of a point lower. We report this protocol because the FERAL^16^ and VideoPrism^33^ comparisons use it, and matching it improves comparability with those reported results. In the three-way protocol, a validation subset was carved out of the training portion only; the test subset was never used during training or checkpoint selection. The checkpoint was selected by validation frame-mAP, and threshold-dependent metrics used cutoffs tuned on validation and applied unchanged to test. For fly courtship, the three-way protocol was run as video-level five-fold cross-validation, with each fold contributing its own training, validation and test subsets. Both protocols split at the coarsest granularity allowed by the dataset—whole recordings wherever every behavior class survived such a split, and clips where it did not—and were designed to keep per-class frame rates as similar as possible across subsets. Unless explicitly stated otherwise, the primary results use the two-way protocol for the single-mouse dataset, CalMS21 and PanAf500; video-level five-fold three-way cross-validation for fly courtship; and the three-way protocol for the DeepEthogram benchmarks. Dataset-specific partitioning is described below.

### Single Mouse Behavior Dataset

To assess V-TRACE on spontaneous single-animal behaviors, we compiled a dataset of self-recorded side-view videos of adult wild-type C57BL/6J mice freely behaving in an open-field arena. Videos were captured from a lateral perspective at 30 frames per second (fps). All behavioral episodes were annotated by a single expert annotator using the V-TRACE graphical annotator, so the labels carry one consistent annotation criterion throughout and no inter-rater reconciliation was applied. The annotation covers four ethologically defined categories: rearing, self-grooming, drinking, and eating. These behaviors span a wide range of durations and display profiles, from brief, repetitive self-grooming to extended eating and drinking. The dataset comprised 13 mice across 34 recordings. After clips spanning animal changeovers were excluded, the annotated dataset contained 505 clips of 768 frames (25.6 s) each and carried 1,417 annotated behavioral events in total.

The primary analyses in **Fig. 2** used a two-way clip-level split of the 505 annotated clips: 80% were assigned to training and 20% to a held-out test portion. The last-epoch checkpoint was reported. For the training-data-reduction analysis in **Fig. 2h**, complete annotated clips were subsampled only from the training portion while the held-out test portion remained unchanged; because subsampling occurred after annotation, this analysis measures sensitivity to training-video quantity rather than annotation effort. A separate three-way train/validation/test analysis was performed for **Extended Data Fig. 1a,b**. Drinking and eating occurred in only a limited subset of recordings, so partitioning by recording could not retain every class across all subsets. Both analyses therefore partitioned the data at the clip level, using multi-label iterative stratification to approximately match per-class frame rates; validation was used for checkpoint selection only in the three-way analysis. Because clips were assigned individually, clips from the same recording and animal could fall in different subsets, so these analyses do not establish generalization to unseen recordings or animals. Model performance was evaluated using mean average precision (mAP) on held-out test clips.

### Caltech Mouse Social Interaction (CalMS21)^32^

The CalMS21 dataset captures social interactions between pairs of freely behaving laboratory mice during a resident–intruder assay, recorded from a top-view camera at 30 fps. The dataset contains frame-level behavioral annotations across three ethologically defined categories: attack, investigation, and mount, reflecting distinct patterns of agonistic and affiliative interactions. The two-way partition is the train–test split of the original challenge’s task 1: 70 training videos and 19 held-out test videos (262,107 test frames). The challenge provides no validation subset, so training runs for the full 10 epochs and the last-epoch checkpoint is the one reported; the backbone comparison additionally disables per-epoch evaluation altogether, so those runs never see a score of any kind before the one reported. The three-way partition keeps the 19 official test videos completely untouched and carves the validation subset out of the 70 training videos only, partitioning them at the video level into 56 training and 14 validation videos. Three such partitions were built as three folds of a balanced five-fold partition of the 70 videos, so the validation folds are mutually disjoint and every video used for validation is held out exactly once across the three runs; each fold was optimized to match the per-class frame rates of the full training set (maximum relative per-class deviation below 0.7%). Their video composition is listed in Supplementary Table 1. Unlabeled frames were treated as a separate “other” background class to enable the model to suppress false detections during inactive periods. Model performance was evaluated using mAP.

### Caltech Fly Courtship Dataset^34^

This dataset contains videos of pairs of *Drosophila melanogaster* engaged in courtship interactions. Videos were recorded from a dorsal view under controlled laboratory conditions at 30 fps, with expert-annotated temporal boundaries for three courtship-specific behaviors: wing extension (the male’s courtship “song” display), circle, and copulation. These behaviors are characterized by brief, rapidly alternating bouts and require precise temporal localization. We merged the *hyperactive* (10 videos) and *wild-type* (21 videos) recordings into a single dataset of 31 videos of 54,000 frames (30 min) each, 1,674,000 frames in total. Two sparse annotation categories were excluded from scoring: “unknown” (0.46% of frames) and “copulation attempt” (0.07% of frames, 104 bouts in the entire dataset, too thin to balance across folds).

This dataset provides no standard partition of any kind, so the partitioning is ours: both reported numbers come from a single scheme, video-level five-fold cross-validation under the three-way protocol. Folds were formed by grouped *k*-fold over recordings (held-out sizes 6/6/6/6/7, against 25/25/25/25/24 training videos), so every clip of a video falls in the same fold and no video is ever split across training and evaluation. Fold assignment was optimized under a hard constraint of exactly two *hyperactive* videos per fold, minimizing the worst-case per-class deviation in both frame prevalence and bout count relative to the full dataset (achieved worst-class deviation: 2.6% for copulation, 4.1% for circle, 5.1% for wing extension). Each fold in turn serves as the held-out test set while the remaining videos are used for training, with the validation subset used for checkpoint selection carved out of those training videos only. Evaluation covers the full timeline of the held-out videos, including all unannotated background frames. Per-fold performance and the mean across the five folds are given in **Extended Data Fig. 1e–g**; the single-fold value quoted in the main figure is the best-performing of the five folds. Model performance was evaluated using mAP.

### Pan African Programme Primate Behavior (PanAf500)

The PanAf500 dataset is a subset of the PanAf20K^35^ collection, comprising 500 camera-trap video clips of wild chimpanzees (Pan troglodytes) recorded across 18 field sites in tropical Africa as part of the Pan African Programme: The Cultured Chimpanzee. The videos capture a wide range of locomotor, postural, and social behaviors under diverse lighting and environmental conditions. Behavioral annotations are provided at the individual level for each clip: every frame carries one bounding box per visible chimpanzee, together with that individual’s within-video identity and behavior label. The annotated behavior categories include walking, standing, sitting, climbing up, climbing down, hanging, running, sitting on back, and camera interaction. PanAf500 is evaluated under the two-way protocol: we keep the 400 training videos of the split that ships with the dataset and merge its 25 validation and 75 test videos into a single 100-video held-out set. This increases the number of videos available for held-out evaluation, which is useful because the rare classes (camera interaction, climbing down) have very few bouts and a 25-video validation subset does not estimate their AP stably. No checkpoint selection is therefore performed and the last-epoch checkpoint is the one reported.

Unlike the laboratory datasets, the chimpanzees in PanAf500 are almost never alone: a single camera-trap frame typically contains several individuals performing different behaviors simultaneously, so the frame as a whole has no single behavior label and whole-frame training forces the model to average over mutually inconsistent targets. We therefore used the dataset’s own individual-level annotations to separate the chimpanzees before classification. Chimpanzee bounding boxes were linked across frames by their annotated within-video identity into per-individual tracks, and each track was converted into its own video by cropping the region of interest spanned by that individual across its lifetime. Each cropped track carries only its own animal’s behavior labels, so behavior recognition is performed per individual rather than per frame; the video-level multi-label target used by the whole-frame control is the pooling of these per-individual labels. Tracks inherit the subset of the video they come from, so the split proportions are unchanged by cropping. Cropping raises mean AP by 11.2 points over the whole-frame control, with gains concentrated in behaviors defined by the animal’s own posture and losses confined to the three behaviors defined by whole-frame context (sitting on back, running, standing; **Extended Data Fig. 3**). Model performance was evaluated using mAP.

### DeepEthogram Benchmarks (Mouse-Homecage and Mouse-Social)

To compare against a published supervised baseline, we evaluated V-TRACE on the two mouse datasets released with DeepEthogram^14^. Mouse-Homecage contains 12 top-view videos annotated with seven behaviors (in hut, object 1 interaction, object 2 interaction, rearing, grooming, interacting with the hut, poking out of hut); Mouse-Social contains 12 videos annotated with five social behaviors (nose-to-nose, nose-to-body, body-to-body, chasing, anogenital sniffing).

Both DeepEthogram benchmarks were evaluated under the three-way protocol only; no two-way partition was constructed for them. For each dataset we built five video-level splits of 7 training, 2 validation, and 3 test videos, following the 60/20/20 fractions of the original DeepEthogram protocol, with every behavior class present in all three subsets of every split; their video composition is given in **Supplementary Table 2**. The checkpoint is selected by validation frame-mAP and the per-class F1 cutoffs are tuned on validation and applied unchanged to the three test videos. The reported score is the mean over the five splits; per-behavior F1 for each split, together with the micro-averaged F1 comparison against DeepEthogram, is given in **Extended Data Fig. 2**.

### Model Architecture

V-TRACE is a frame-resolved temporal behavior-detection framework that predicts a behavior probability for every frame of an untrimmed video. The pipeline consists of three components: a frozen spatiotemporal feature backbone with lightweight trainable adapters, a multi-scale temporal projection and a temporal prediction head. The configurations evaluated in this study use dense per-frame classification heads, with contiguous runs of frame-wise predictions defining the onset and offset boundaries of behavioral bouts.

### Backbone: frozen video transformer with trainable temporal adapters

Spatiotemporal features are extracted by a VideoMAEv2^24^ ViT-B encoder (embedding dimension 768, 12 transformer layers, 12 attention heads, ∼86M parameters), initialized from a publicly released checkpoint pretrained by its original authors using masked autoencoding on Kinetics-400. We did not perform the large-scale pretraining of the VideoMAEv2 or V-JEPA2 backbones used in this study. The backbone weights are kept entirely frozen; the only trainable backbone components are lightweight temporal adapter modules inserted at every transformer layer (4.0M parameters, ∼4.6% of the backbone). Input frames are resized according to the input-resolution requirement of each backbone. For the default VideoMAEv2 ViT-B configuration, frames are embedded with a 16 × 16 spatial patch size and a temporal tubelet of 2 frames. Long videos are processed as windows of 768 frames (25.6 s at 30 fps): each window is divided into 48 non-overlapping 16-frame micro-clips that are independently encoded. With a temporal tubelet of 2 frames, spatial pooling yields eight temporal positions per micro-clip; concatenating the 48 chunks therefore produces a 384-step temporal sequence. This sequence is linearly interpolated to 768 steps to yield one 768-dimensional feature vector per frame. Positional information within each chunk is supplied by the pretrained encoder, and absolute sinusoidal positional encoding is applied to the concatenated temporal sequence before multi-scale temporal modeling.

### Dense frame classification head

The per-frame feature sequence is projected to 512 channels and expanded into a multi-scale temporal pyramid. The default model uses a temporal convolutional network (TCN) classification head to map this temporal representation to per-frame class probabilities over all behavior categories plus an explicit background channel, trained with a softmax (single-label) objective. The TCN uses residual dilated convolutions with dilation doubling from 1 to 32, providing a receptive field of approximately 127 frames. For the classification-head configurations evaluated here, no non-maximum suppression is applied; predictions from overlapping sliding windows are averaged, and contiguous runs of the arg-max class form the predicted ethogram.

### Per-Frame Head Comparison

Because the model applies a per-frame classifier to a shared feature sequence, the temporal head is a modular component that can be varied while holding the remaining pipeline fixed. We therefore made the head pluggable and compared six temporal-head families on the same features. All heads share one interface — they map a channel-by-time feature sequence to per-frame class logits at the same temporal resolution — and are selected by a single configuration key, so switching between them changes nothing else in the pipeline.

Six heads were compared. A **convolutional** stack (kernel 3, receptive field about 5 frames, 1.6M parameters) provides a local convolutional baseline with a fixed small context. A **temporal convolutional network** in the MS-TCN style^38^ (residual dilated convolutions with the dilation doubling from 1 to 32, receptive field about 127 frames, 6.6M) keeps locality but reaches much further in time. A **temporal transformer^39^ (global scaled dot-product attention with sinusoidal position encoding, 6.6M) replaces locality with content-adaptive global relations. An ASFormer-style head^40^ (6.6M) restricts that attention to a band whose window doubles with depth, giving content-adaptive local relations as an intermediate case. A state-space head^41^ (a diagonal S4D-style linear recurrence realized as a parallel causal convolution, 7.6M) models long-range dependence by recurrence rather than attention. A dynamic feature-aggregation** head after DyFADet^42^ (2.4M) aggregates over a local window with input-dependent kernels. In addition, **TriDetHead^23^** is another optional boundary-regression architecture in V-TRACE for applications that require direct regression of temporal boundaries.

How a head attaches to the backbone was treated as part of its design rather than as a free parameter. Each head was connected at the temporal scale or scales appropriate to its published architecture, avoiding a forced common attachment that could favor or disadvantage particular head families. The comparison holds everything but the head fixed: the same frozen VideoMAEv2 ViT-B backbone with the same trainable adapters, the same schedule, optimizer, augmentation and loss, one seed, and the same CalMS21 dataset. Heads are ranked by frame-mAP on the held-out clips, and per-class AP is reported alongside the mean, because the question is not only which head is best on average but whether different behaviors prefer different biases. To separate the effect of the head from that of the features it is measured on, the comparison was repeated across pretrained backbones: **Fig. 5g** reports frame-mAP for every combination of six pretrained video encoders (V-JEPA2^28^, MAEv2-L (K400), MAEv2-B (MABe), MAEv2-B (K710), MAEv2-B (K400) and MAEv2a-S (K710)) and six temporal heads (TCN, DyFADet, Attn, Conv, ASFormer and SSM), so that reading down a column isolates the contribution of the backbone and reading across a row shows whether the head ordering is stable.

### Feature Distillation

The adapters are initialized by feature distillation^43,44^ from a larger teacher before any supervised training.

The teacher is a V-JEPA2^28^ ViT-L encoder fine-tuned on the training videos of the corresponding dataset; the student is the frozen VideoMAEv2^24^ ViT-B described above, with only its adapters (plus a temporary 768→1024 projection) trainable. For 750 optimization steps (AdamW, learning rate 1 × 10⁻^4^, weight decay 0.05, gradient clipping at 1.0, batch size 1), the student is trained to match the teacher’s final feature sequence on unlabeled training-set windows, under a cosine-similarity loss plus 0.5 × mean-squared error. The projection is then discarded and the distilled adapters initialize the downstream model.

Dataset-specific teachers were used for the mouse datasets (CalMS21, single-mouse, and the DeepEthogram benchmarks); the fly courtship model reuses the CalMS21-distilled adapters without any fly-specific distillation.

The feature-matching stage uses only unlabeled video from the training split and never uses test video. On CalMS21, feature distillation increased frame-mAP by 3.39 percentage points (91.54% without distillation versus 94.93% with distillation; **Fig. 5d**) without changing the deployed student architecture, because the teacher is discarded after the 750 distillation steps. The fly courtship model reused the CalMS21-distilled adapters without fly-specific distillation, providing an example of reuse of the same initialization in a different species and recording setting. As with other transferred representations, the resulting initialization may retain biases present in the teacher features.

### Behavior-Targeted Window Sampling

Training windows are drawn by a behavior-targeted sampler that combines full coverage with oversampling of rare behaviors. A sparse base pass tiles every training video with non-overlapping 768-frame windows (position-jittered by up to 12.5% of the window per epoch), preserving the dataset’s natural class and background distribution. On top of the base pass, every bout of a *rare* class receives additional anchor windows whose positions are re-drawn every epoch from strata spanning the bout’s possible phases within the window, from “just entering at the window tail” to “just leaving at the head”, so that across epochs the model sees each rare bout at a continuum of temporal offsets. A class is deemed rare relative to its own dataset: when its share of the timeline falls below one half of the mean class prevalence, or below one quarter of the most frequent class’s share. The number of anchors per bout scales with the inverse square root of the class’s labeled-frame count (2 to 12 anchors per bout), so that a class with few long bouts and a class with many short bouts receive comparable additional exposure. Validation and test data are never resampled: evaluation always slides uniformly over the full video.

### Training

All models are trained with AdamW (weight decay 0.025) with a 2-epoch linear warmup followed by cosine annealing (1-epoch warmup for fly courtship). The training length is matched to dataset size: 10 epochs for CalMS21, PanAf500 and the single-mouse dataset, 5 for fly courtship, and 20 for the DeepEthogram splits. The frozen ViT receives no gradient; adapter parameters use a learning rate of 1 × 10⁻^4^ (weight decay 0.05) and all remaining components 7 × 10⁻^5^. The batch size is 2 (4 for the DeepEthogram splits). Training uses bfloat16 automatic mixed precision, gradient clipping (max norm 1), and an exponential moving average of model weights, which is also the weight set used at evaluation. Data augmentation comprises random temporal speed perturbation (0.7–1.3×), horizontal flipping (p = 0.5), random rotation (up to 180°, p = 0.8), and TrivialAugment image transformations; frames are normalized with ImageNet statistics, matching the preprocessing used by the pretrained VideoMAEv2 checkpoints. Under the three-way protocol the checkpoint with the highest validation frame-mAP is selected; under the two-way protocol, which has no validation subset—the single-mouse dataset, CalMS21 and PanAf500—no checkpoint selection is performed and the last-epoch checkpoint is reported (see Datasets). For the primary 80/20 single-mouse analysis in **Fig. 2**, the last-epoch checkpoint is reported; the separate three-way analysis in **Extended Data Fig. 1a,b** uses validation-based checkpoint selection. The full recipe trains and runs inference on a single consumer GPU: peak training memory is 12.2 GiB at batch size 2, within the 16 GB of a GeForce RTX 5070 Ti.

### Loss Functions

The model is trained with a single dense classification loss: per-frame cross-entropy over the behavior classes and the background channel, with label smoothing of 0.1 and mixup (α = 0.8). Class imbalance is handled by inverse-square-root frequency class weights, so rare behaviors contribute larger per-frame gradients without altering the sampler’s coverage guarantees. Frames outside any annotated episode supervise the background channel explicitly, which lets the model suppress false detections during inactive periods rather than relying on score thresholds alone.

### Evaluation

Model performance is evaluated with frame-level mean Average Precision (frame-mAP): for each behavior class, per-frame one-vs-rest average precision is computed over every frame of the held-out videos, with background frames as negatives, and the mean is taken over the foreground classes; the background class itself is excluded from the average. Frame-mAP is threshold-free and weights errors by their duration, matching how ethograms are consumed downstream. For threshold-dependent summaries (precision, recall, F1), per-class score cutoffs are tuned on the validation split and applied unchanged to the test split. At test time, videos are processed with 768-frame sliding windows at 50% overlap and window predictions are averaged.

### Statistical Analysis

To compare behavioral profiles, eight recording sessions per group were acquired from wild-type (WT, n = 3 mice) and Alzheimer’s disease model (AD, n = 3 mice) animals. V-TRACE-generated ethograms were used to quantify the percentage of recording time spent in each behavioral category (rearing, self-grooming, drinking, and eating) per recording session. Each recording session was treated as an observational unit. Group differences were assessed using two-sided Mann–Whitney U tests (scipy.stats.mannwhitneyu) for each behavior independently. Exact two-sided *p* values are reported in the figure legends and in the Results rather than significance stars, and no corrections for multiple comparisons were applied. Cumulative behavior time curves are plotted as the mean across videos within each group, with shading indicating the standard error of the mean (s.e.m.).

## Data Availability

The single-mouse videos and annotations generated for this study are available from the corresponding authors upon reasonable request. The public datasets analyzed in this study are available from their original repositories: CalMS21 (https://data.caltech.edu/records/s0vdx-0k302), Fly-vs-Fly Courtship (https://data.caltech.edu/records/zrznw-w7386), PanAf500/PanAf20K (https://data.bris.ac.uk/data/dataset/1h73erszj3ckn2qjwm4sqmr2wt), and the Mouse-Homecage and Mouse-Social datasets distributed with DeepEthogram (https://github.com/jbohnslav/deepethogram).

## Code Availability

The graphical user interface and source code for V-TRACE, including training, distillation, and inference scripts, together with configurations for the released models, are publicly available on GitHub: https://github.com/KunmingS/TRACE. V-TRACE is distributed as a locally installed program, and users launch its graphical interface on their computer. Model checkpoints are hosted on the repository’s release page and can be downloaded through the installed package. The repository also provides model-download support, Jupyter and Colab notebooks, a demonstration package containing an example video, a V-TRACE-compatible annotation file and a trained model, and utilities for converting public datasets, including CalMS21. User-reported issues are tracked through the public repository, with maintenance and release oversight by the author team and corresponding authors. Claude Code was used to assist with coding during V-TRACE software development. The software is distributed under the Apache License 2.0.

