## Supplementary Figures for "V-TRACE: End-to-end temporal inference and annotation of animal behaviors from video"

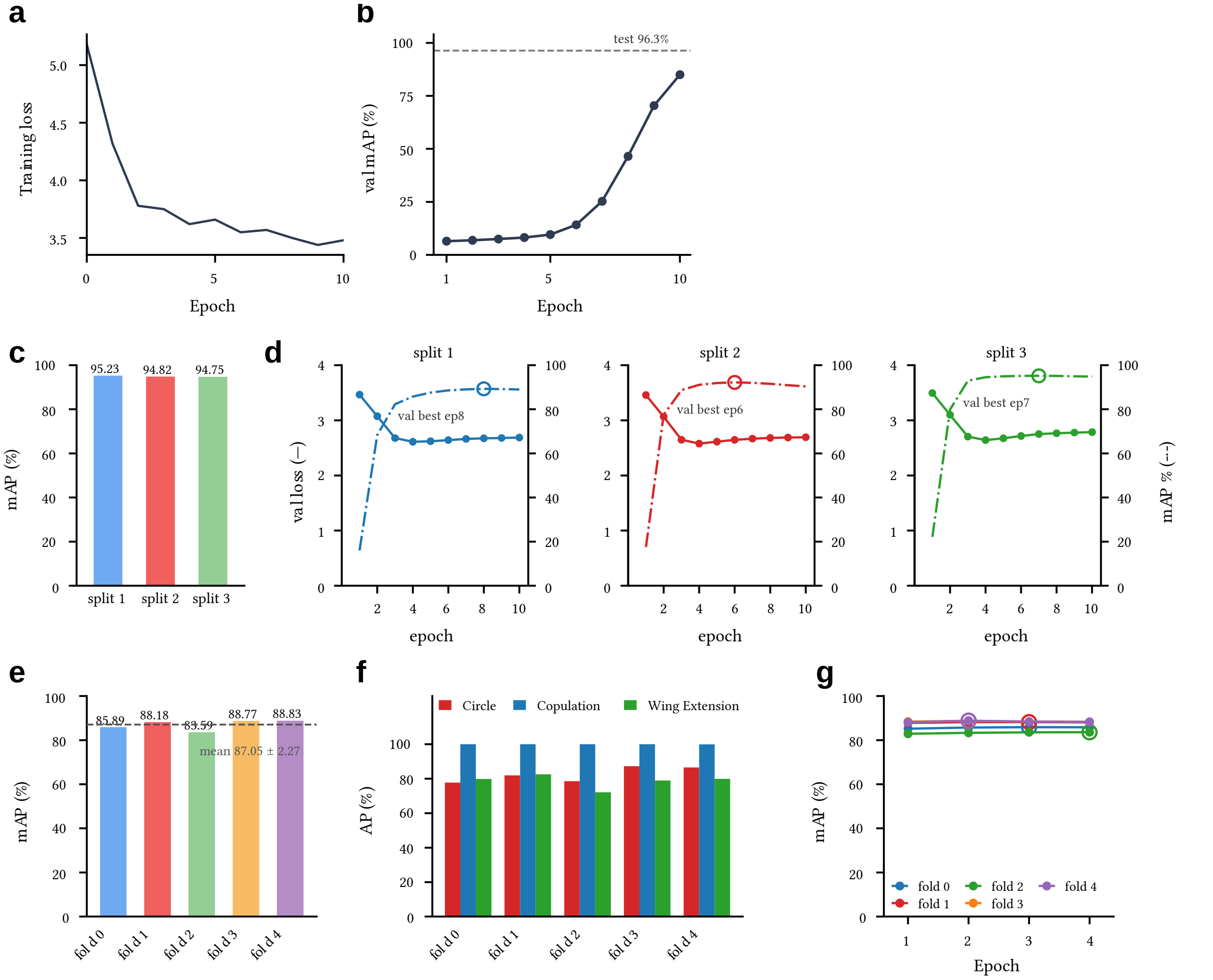


**Extended Data Figure 1 | V-TRACE performance across validation partitions and video-level folds. a,b,** Training loss (a) and validation frame-mAP (b) of the distilled MAEv2-B configuration on the single-mouse open-field dataset using a three-way train/validation/test split. The dashed line in b indicates the corresponding held-out test frame-mAP. **c,** Test frame-mAP of the distilled MAEv2-B configuration across three independent train/validation partitions drawn from the official 70-video CalMS21 training set; each selected checkpoint was evaluated on the same 19-video official test set. **d,** Validation loss and validation frame-mAP over training epochs for the corresponding CalMS21 partitions; open circles indicate the selected checkpoints. **e-g,** Performance across five video-level folds of the Fly-vs-Fly Courtship dataset, showing held-out frame-mAP (e), per-behavior AP (f) and validation frame-mAP during training (g); open circles in g indicate the selected checkpoints.

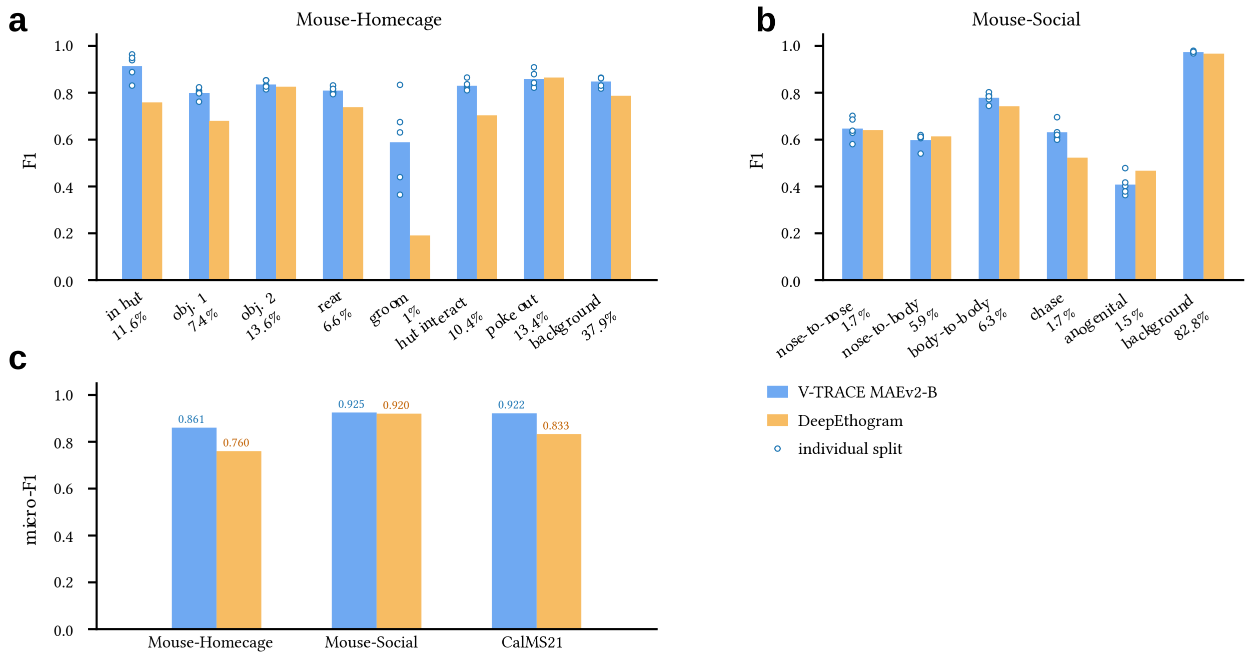


**Extended Data Figure 2 | V-TRACE performance on the DeepEthogram benchmark datasets. a,b,** Per-behavior F1 scores for V-TRACE (MAEv2-B) and DeepEthogram on Mouse-Homecage (a) and Mouse-Social (b). V-TRACE bars show the mean across five independent video-level splits, with individual splits indicated by open circles. Percentages beneath the behavior labels indicate class prevalence. **c,** Micro-averaged F1 for V-TRACE and DeepEthogram on Mouse-Homecage, Mouse-Social and CalMS21. The DeepEthogram values for Mouse-Homecage and Mouse-Social are published results; the CalMS21 value was obtained by evaluating the full DeepEthogram pipeline under the corresponding held-out protocol.

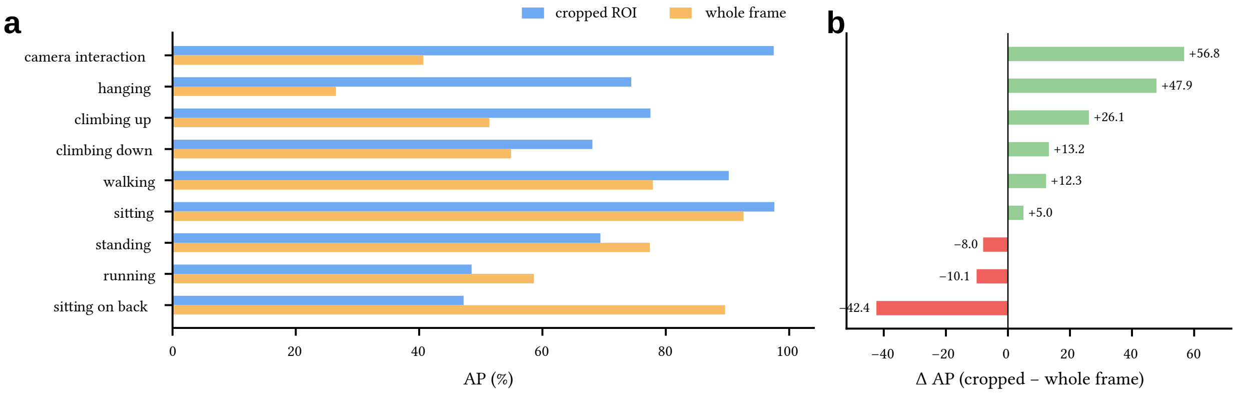


**Extended Data Figure 3 | Subject-centered cropping differentially affects behavior classification on PanAf500. a,** Per-class AP for V-TRACE models trained using subject-centered chimpanzee regions of interest (cropped ROIs) or whole-frame inputs. **b,** Difference in per-class AP between cropped-ROI and whole-frame inputs. Positive values indicate higher AP with cropping and negative values indicate lower AP. Cropping increased AP for six of nine behaviors but reduced AP for sitting on back, running and standing.

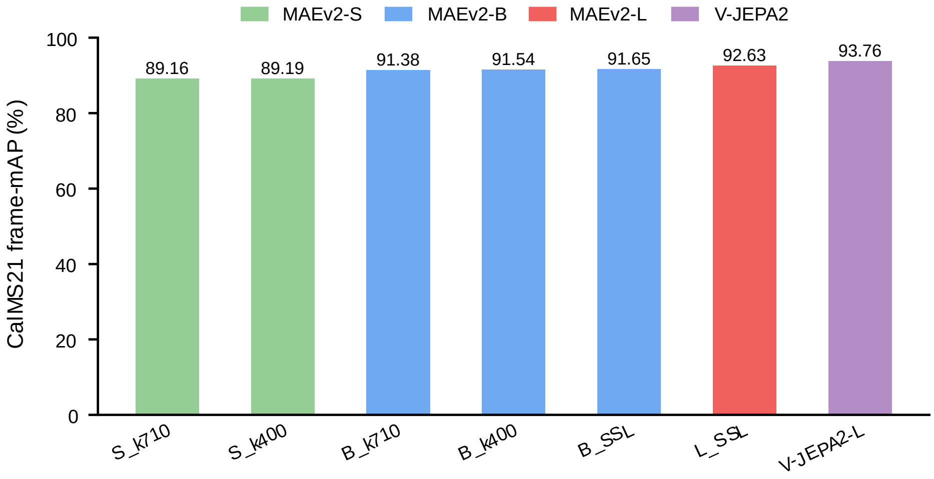


**Extended Data Figure 4 | Benchmarking backbone architecture, scale and pretraining under full fine-tuning on CalMS21.** Test frame-mAP for seven fully fine-tuned encoder configurations on CalMS21. Each configuration was trained on the official 70-video training set and evaluated on the 19-video official test set in one run, using the fixed epoch-9 checkpoint after 10 epochs without validation-based checkpoint selection. For the model labels, S, B and L indicate small, base and large model scales, respectively; K710, K400 and SSL denote Kinetics-710, Kinetics-400 and self-supervised learning, respectively. All runs used the same data partition, epoch count and checkpoint-selection protocol.

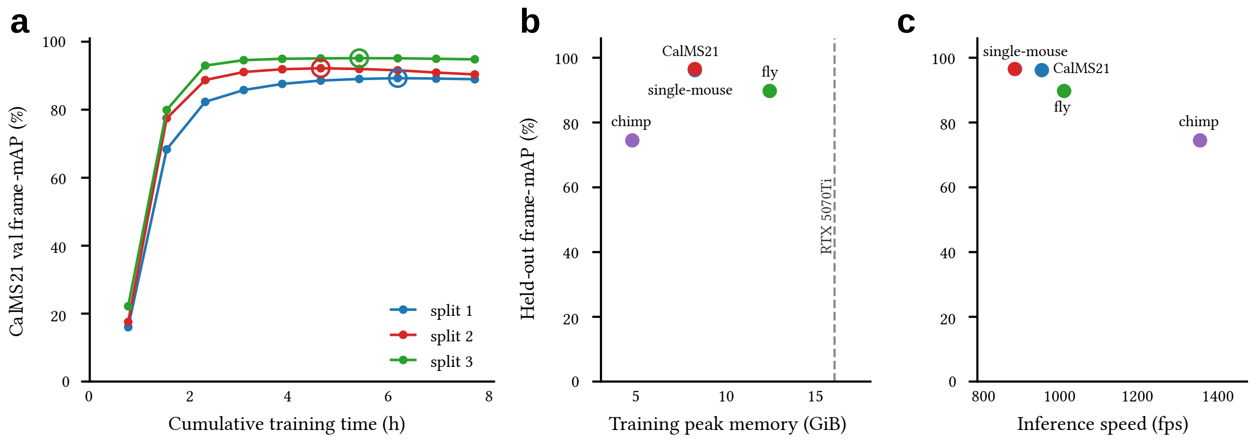


**Extended Data Figure 5 | Training and inference of the default V-TRACE configuration on a 16-GB GeForce RTX 5070 Ti. a,** CalMS21 validation frame-mAP as a function of cumulative training time across three validation partitions; open circles indicate the selected checkpoints. **b,** Held-out frame-mAP versus peak training GPU memory for the single-mouse, CalMS21, fly courtship and PanAf500 datasets. The dashed line indicates the 16-GiB GPU-memory capacity. **c,** Held-out frame-mAP versus end-to-end inference throughput for the same datasets. Each point in b and c represents one dataset; 'chimp' denotes PanAf500. The default configuration uses the distilled MAEv2-B encoder with the TCN temporal head.
